# Effects of dibutyl phthalate (DBP) on life history traits and population dynamics of *Daphnia magna*: comparison of two exposure regimes

**DOI:** 10.1101/2025.03.05.641620

**Authors:** Jérémie Ohanessian, Sophie Martine Prud’homme, Ellis Franklin, Géraldine Kitzinger, Catherine Lorgeoux, Elise Billoir, Vincent Felten

## Abstract

Phthalates are chemical products used as additives in the synthesis of plastics to increase their flexibility and resistance. Among the most frequently detected phthalates in the environment is dibutyl phthalate (DBP). Although ubiquitous in freshwater environments at concentrations in the ng/L to µg/L range, studies on the effects of DBP on *Daphnia magna* have overlooked the potential effects of exposure at low concentrations (µg/L order) and without solvent. Our work focuses on the chronic effects of DBP on life history traits (survival, growth, reproduction) in the freshwater model species, *D. magna* (Crustacea). Organisms were exposed to two exposure regimes: from the beginning of embryonic development or from the adult stage, at the third brood, when most of the energy investment in growth is complete.

The results obtained show that the DBP exposure regime is an essential parameter in the effects on *D. magna* life history traits. While no significant response was observed in organisms exposed at the adult stage, disturbances to survival, growth and reproduction were observed at concentrations as low as the µg/L range in organisms exposed at the beginning of embryonic development. The results also demonstrated that exposure to a concentration gradient of DBP results in different dose-dependent response typologies, depending on the life history traits measured. For survival, the dose-response relationship was non-monotonic, with greater juvenile mortality at intermediate concentrations (100, 280 and 500 µg/L) than at higher concentrations (1000 and 2000 µg/L). For effects on growth and reproduction, the responses followed classic monotonic dose-response relationships with low sensitivities. At the end of the experiment, the EC10-25d values were 1.01 µg/L for reproduction and 79.22 µg/L for growth. However, when calculated at earlier timepoints, the EC10-4d and the EC10-7d for growth were 0.25 µg/L and 1.16 µg/L. Furthermore, projecting these results to the population level suggests that exposure to DBP from early embryonic development leads to a decrease in growth rate and a change in population structure.

## 1. Introduction

Phthalates are synthetic esters of phthalic acids, used as additives in the synthesis of plastics polymers such as polyvinyl chloride (PVC), polyethylene (PET) and polycarbonate (PC) to increase their flexibility and resistance. Due to their properties, they are also used in various applications, such as enhancing the adhesive effects of pigments and preserving perfumes. As a result, they are used in numerous commercial and consumer products such as adhesives and sealants, paints and coatings, inks, cosmetics, floor coverings, furniture, lubricants, fabric, textiles and leather (Net *et al*., 2015). First produced in 1920, their worldwide production has continued to grow, reaching millions of tons in the 2010s (Net *et al*., 2015). Direct industrial discharges, leachate from materials during use, recycling, and disposal, as well as from legacy plastics, lead to an ubiquitous contamination of global environments by phthalates, including air, soils and surface waters (Net *et al*., 2015; Gao & Wen, 2016; Cao *et al*., 2022). Among the most frequently detected phthalates in the environment is dibutyl phthalate (DBP). Despite DBP being restricted for specific uses by several countries (e.g. forbidden in food packages in EU; Net *et al*., 2015), it is still one of the most commonly found and abundant phthalic compounds in surface freshwaters (Net *et al*., 2015; Gao & Wen, 2016). DBP concentrations in surface freshwater is commonly measured at concentrations ranging from ng/L to tens of μg/L (Shi *et al*., 2016; Paluselli *et al*., 2018; Nantaba *et al*., 2021), with higher concentrations (hundreds of µg/L) occurring in sites undergoing high industrialisation (Wu *et al*., 2013; Dada & Iken, 2018). Exposure to DBP is thus nearly unavoidable for aquatic wildlife. The high production and use of phthalate-containing products, along with the projected increase in plastic production in the coming decades (Geyer *et al*., 2017) suggest that this situation will persist, if not worsen.

DBP is described as an endocrine disruptor able to cause reproductive toxicity in mammals, especially males (Radke *et al*., 2018; Arzuaga *et al*., 2020; Czubacka *et al*., 2021) and is classified as reprotoxic by the European CHemicals Agency (ECHA). However, there is limited ecotoxicological data testing for DBP effects on aquatic invertebrates, even for the microcrustacean *Daphnia magna*, which is a widely used primary consumer model organism in risk assessment. The few studies available suffer from several weaknesses limiting their ability to inform the DBP ecotoxicity on *D. magna.* (1) Most of them concern only acute exposure to DBP (<48h) (Adams *et al*., 1995; G. L. Huang *et al*., 1999; Jonsson & Baun, 2003; B. Huang *et al*., 2016a; Wei *et al*., 2018a; Shen *et al*., 2019a), and to our knowledge, only six studies investigated the chronic toxicity of DBP on *D. magna* (McCarthy & Whitmore, 1985a; Defoe et al., 1990a; Rhodes *et al.,* 1995; Wei *et al*., 2018b; Seyoum & Pradhan, 2019a; Jin *et al*., 2024). All those with chronic DBP exposure, covering exposure times from 16 to 60 days, begin on neonate individuals within 24 hours after hatching, thus excluding the embryonic development period from exposure. However, aquatic organism populations are continuously exposed to DBP throughout their life cycle, and embryonic development is a particularly sensitive period to pollutants. Several studies have highlighted the need to study the different life stages to fully understand the effects of pollutants (Abe *et al*., 2001; Sobral *et al*., 2001; Palma *et al*., 2009; Ton *et al*., 2012). The characterisation of DBP ecotoxicity on *D. magna* should thus consider a realistic exposure scenario, including all the life cycle phases of individuals, from the beginning of embryonic development to reproduction. (2) The six chronic studies aiming at characterising the effects of DBP on *D. magna* considered only high environmental concentrations (> 50 µg/L) and ignored low to medium contamination levels, limiting their ability to represent the diversity of environmental situations in which populations are exposed. (3) It is also noteworthy that, except for the study by Rhodes *et al*. (1995), the other studies used either DMSO or acetone as DBP solvents at various final concentrations in the exposure media. Yet, the use of solvents can be a confounding factor in ecotoxicological studies, and where technically possible, no solvent should be used (Green & Wheeler, 2013; Hutchinson *et al*., 2006). Given DBP’s solubility in water (11,2 mg/L, (Staples *et al*., 1997)), it seems unnecessary to use additional solvents to investigate its ecotoxicity at environmentally realistic concentrations. (4) Furthermore, apart from Rhodes *et al*. (1995), none of these studies mentioned any strategies to limit external contamination of experimental units, and DBP background contamination in control media is not always measured (Seyoum & Pradhan, 2019a; Jin *et al*., 2024). Nevertheless, considering the leaching of DBP from usual lab consumables and equipment, it is of utmost importance to address this issue (Reid *et al*., 2007a).

Linking the effects of toxic substances on individual performance to their consequences at the population level is a major aspect of ecotoxicology, crucial for improving its environmental relevance (Forbes *et al*., 2011). Population matrix models can be developed from life cycle experiments to translate the effects of exposure on individual level parameters into population level metrics (Leslie, 1948; Caswell, 2001). In this context, population dynamics models have proven very useful for assessing the demographic consequences of alterations in life history traits in chronic exposure models (Raimondo *et al*., 2009; Biron *et al*., 2012; Prud’homme *et al*., 2017). However, to date, no study investigated the population-level consequences of DBP exposure in *D. magna* nor any other invertebrate.

In this context, our objectives were (1) to characterise the impact of DBP exposure on *D. magna* life history traits (survival, growth and reproduction) across a wide range of nine concentrations, from low environmental concentrations (0.5 µg/L) to lethal ones (2000 µg/L; Rhodes *et al*., 1995), without using additional solvents, and (2) to project the consequences of altered life history traits at the population-level. The tested concentration gradient included environmental concentrations (0.5, 1 and 10 μg/L), the NOEC of DBP for chronic exposure in invertebrates according to ECHA (100 μg/L), concentrations at which disruption of lipid storage in *D. magna* has been shown (280 μg/L; Seyoum & Pradhan, 2019a), and induction of oxidative stress after acute exposure (500 μg/L; Shen *et al*., 2019a). The two higher concentrations, 1000 and 2000 μg/L, were included to extend the range of exposure with concentrations likely leading to effects on life history traits. Throughout the experimentation, we paid specific attention to limiting external phthalic compound contamination in our exposure media by avoiding the use of plastic consumables. Two DBP exposure regimes were considered: a full life exposure, covering the embryonic development to the reproductive phase, and a partial adult exposure covering only the reproductive phase, when most of the energy investment in growth is complete. Our work provides novel insights into the effects of DBP on *Daphnia magna* and its potential consequences on population dynamics.

## 2. Materials and Methods

### 2.1. *D. magna* origin and acclimation

*Daphnia magna* clone A (or clone 5; Felten *et al*., 2020) were supplied by the LIEC laboratory (Metz, France). Since 1974, this population/clone of *D. magna* has been maintained in a parthenogenetic phase of reproduction according to ISO 6341 (1996). To avoid any external uncontrolled contamination of *D. magna* culture media by phthalates or any other plasticiser leaching from plastic laboratory consumables (Reid *et al*., 2007b), all media and food preparation, as well as distribution, were performed exclusively using glassware throughout the experiment. Juvenile individuals from the LIEC laboratory population were isolated within 12 hours after hatching, and were placed in plastic-free experimental conditions to acclimate the daphnids before the beginning of the experiment (Figure 1, “Acclimated generation”).

**Figure 1.**
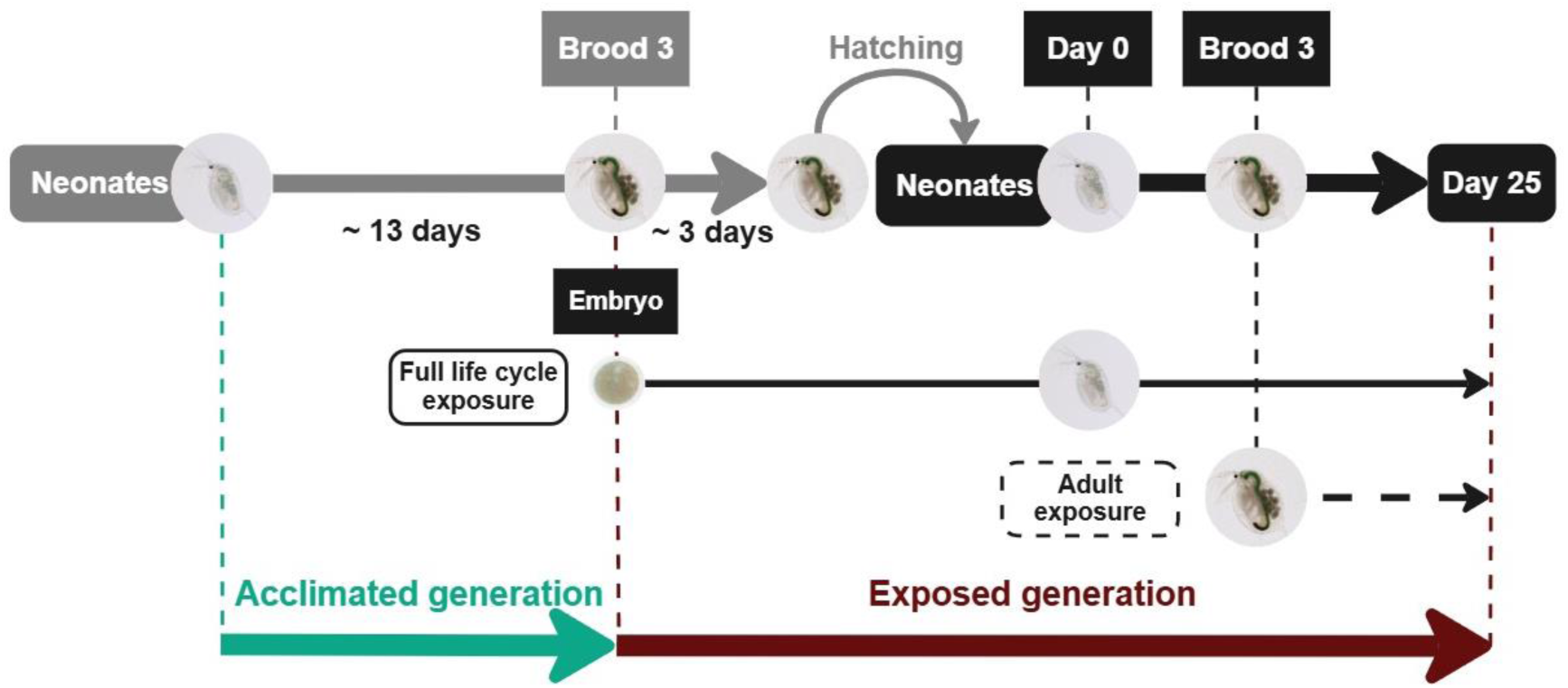
Experimental design for evaluating the impact of DBP on *D. magna* life history traits. Green denotes the first generation acclimated to our phthalate-free experimental conditions. Red denotes the generation exposed to DBP without solvent according to two exposure regimes: (1) full life cycle exposure, starting from the beginning of embryonic development (8 DBP concentrations), and (2) adult exposure, from the third brood (6 DBP concentrations).

These juveniles were placed in 4 batches of 30 individuals at a density of 1 individual per 50 mL in medium M4 at pH 8 ± 0.1 (mean ± SD) (Elendt & Bias, 1990) with an oxygen concentration of 6.58 ± 0.18 mg/L. The medium was renewed twice a week. Daphnids were maintained in a climate-controlled chamber at 21 ± 0.2 °C (Cooled incubator MIR-554-PE), with a light intensity of 800 to 1000 lux and a photoperiod of 16 hours of light and 8 hours of darkness. All individuals in this study were fed daily with a mixture of 50% *Raphidocelis supcatata* and 50% *Chlorella vulgaris* at 100 µg of carbon per day per organism. These unicellular green algae were cultured in plastic-free conditions at 22°C in Oligo LC medium (Graff *et al*., 2003) under light with bubbling. At the end of the culture, the algae were centrifuged and resuspended in M4-pH8 for daphnids feeding.

### 2.2. Daphnids exposure

Acclimated daphnids were used to set up the DBP exposure experiment. During this experiment, the impact of two DBP exposure regimes on *D. magna* life history traits were compared: (1) the full life cycle exposure regime, with an exposure from the beginning of the embryonic development (stage 1) (Toyota *et al*., 2016) until day 25 (Figure 1, “full life cycle exposure”) and (2) the adult exposure regime, with an exposure from brood 3 (day 14 of the experiment), when most of the energy investment in growth had been completed, until the end of the experiment on day 25 (Figure 1, “adult exposure”) corresponding to the beginning of the seventh brood in control daphnids. For the full life cycle exposure regime, daphnids were exposed to 8 nominal concentrations of DBP (0.5, 1, 10, 100, 280, 500, 1000 and 2000 µg/L) whereas exposure was limited to the 6 lowest DBP nominal concentrations (0.5 to 500 µg/L) for the adult exposure regime. Real DBP concentrations were measured (mean ± S.D.) at: control: 0.14 ± 0.02 µg/L, 100 µg/L: 93.6 ± 29.2 µg/L, 280 µg/L: 225.1 ± 14.4 µg/L, 500 µg/L: 419.4 ± 43.5 µg/L, 1000 µg/L: 876.3 ± 130.5 µg/L, 2000 µg/L: 1695 ± 182.7 µg/L. Media contaminated at 0.5, 1 and 10 µg/L could not be measured due to too limited available volumes at these low concentrations.

#### 2.2.1 Exposure media contamination and control

Dibutyl phthalate (DBP - CAS No. 84-74-2) was purchased, with reported purity greater than 99% (Sigma-Aldrich). A solvent-free DBP stock solution was prepared at the beginning of the experiment at a nominal concentration of 10 mg/L in M4 medium-pH8 (real DBP concentration: 9.6 ± 1.17 mg/L). As this concentration is below the solubility limit of DBP in water of 11.2 mg/L (PubChem, 2022), no additional solvent was required for stock and exposure solutions.

DBP was added to M4 medium-pH8 with a glass pipette and agitated in darkness for 72h using a glass magnetic stir bar. The effective DBP concentration of this stock solution was measured before the beginning of the experiment (three technical replicates), using an Ultimate 3000 Rs chromatography system with UV detection at 230 nm (DAD) (Dionex, CA, USA). Quantification was performed on an injected volume of 10 µL, using an Acquity UPLC HSS T3 2.5 µm, 2.1×100 mm column (Waters, Germany) as stationary phase, and 60% acetonitrile and 40% H_2_O as mobile phase, under a flow of 8 ml/min at 25°C. During all the experiment, the DBP stock solution was stored in a cold room (around 4°C) and obscurity.

Exposure media were renewed twice a week by serial dilution from the DBP stock solution in M4 medium. Evolution of DBP concentrations in exposure media throughout the experiment was monitored on days 3 to 6 and 17 to 20, through sampling of exposure media immediately after medium renewal (0h), between two renewals (36h) and immediately before the next renewal (72h).

#### 2.2.2 DBP extraction and quantification

DBP was extracted from 100 mL of media sample. Samples were first prepared for extraction by volumisation, dilution (if necessary), and the addition of dDBP, a deuterated surrogate ([2H4]di(n)butylphthalate, supplied by CIL Cluzeau), to assess the extraction recovery for each sample and correct the analytical result. Extractions were performed by Solid Phase Extraction (SPE) with Oasis HLB cartridges (200 mg, 6 mL, Waters), using a Dionex AutoTrace 280 unit (ThermoFisher Scientific). SPE cartridges were first conditioned with hexane, isopropanol, and deionized water. Samples were extracted at a flow rate of 10 mL/min. The SPE cartridges were then dried under nitrogen, rinsed with 3 mL of hexane, and eluted with 5 mL of hexane. Eluted extracts were concentrated to 100 µL under vacuum, after addition of an internal standard dDPP ([2H4]di(n)propylphthalate supplied by CIL Cluzeau).

Analyses were performed on Shimadzu GCMS-QP2010 Ultra and when necessary to increase the sensitivity the Agilent Technologies 7890 GC coupled to a TQ 7010 mass spectrometer. The two instruments were equipped with a silica DB5-MS column (60 m × 0.25 mm id × 0.25 μm film thickness). For these two instruments, an internal calibration was carried out for DBP and dDBP using dDPP as internal standard.

The oven temperature was programmed as follows: 50°C for 1 minute, then increased to 310°C at 20°C/min and held for 10 minutes. The carrier gas was helium at 1.2 mL/min constant flow. The injection was set in splitless mode at 320°C. The GC-MS transfer line temperature was 300°C. Molecule ionization was performed in electronic impact mode at 70 eV. With the triple quadrupole analyser, compounds were followed in multiple reaction monitoring (MRM) to increase the sensitivity of the detection and with the single quadrupole analyser the Single Ion Monitoring Mode was used.

The calibration range was checked with a control standard analysis every 10 samples, and a deviation of less than 20% was accepted. Experimental and analytical blanks were also monitored regularly to assess external contamination. The surrogate extraction efficiency was determined for each SPE extraction and each analytical result was corrected accordingly.

#### 2.2.3 Exposed generation and DBP impact assessment

Synchronous acclimated daphnids (see 2.1), which had just hatched the juveniles of their second brood and just after the third brood (egg deposition into the maternal brood pouch), were grouped into 8 batches of 8 individuals each and 2 batches of 28 individuals each (density of 1 individual per 50 mL). Each batch of 8 individuals was placed in contaminated media (1 per concentration, from 0.5 to 2000 µg/L) to begin the exposure of their offspring from the beginning of stage 1 of embryonic development (Toyota *et al*., 2016) (Figure 1, “exposed generation”, full life cycle exposure regime). At the same time, the 2 batches of 28 individuals were put in phthalate-free media to produce unexposed neonates for the control condition and the “adult exposure” regime (Figure 1, “exposed generation”, adult exposure regime).

Synchronous newly hatched (within 12 hours) juveniles of the exposed generation were individually placed in their test condition (concentration and exposure regime) to initiate a semi-static observation for 25 days inspired by the OECD bioassay 211 (*Test No. 211*, 2012). Eleven daphnids per condition were individually placed in 50 mL of medium M4 at pH 8 (±0.1) (Elendt & Bias, 1990) with the same temperature, light, food and oxygen conditions as described for the acclimated generation. All media were renewed twice a week.

All individuals were monitored daily from day 0 of the experiment (juvenile hatching) until day 25, for survival, egg deposition into brood-pouch, and juvenile hatching. Daphnids were considered dead if they did not move for 15 seconds, even after gentle mechanical stimulation, and were then removed. When hatching occurred, juveniles were counted and immediately removed. Growth of individuals was monitored by measuring the length of *D. magna* at 0, 2, 4, 7, 13 and 25 days for individuals of the full life cycle exposure regime and at 0, 13 and 25 days for those of the adult exposure regime. Pictures of each individual were taken on a digital microscope (KEYENCEVMX-7000) and body length was measured using the “VHX-6000_950F Measurement Data Tabulation Tool” image processing software (KEYENCE, France), from the eye to the base of the caudal spine (Agatz *et al*., 2015; Liu *et al*., 2019). Pictured individuals were placed in a medium drop and immediately put back in their culture media after picture acquisition. The whole process took no longer than 30 seconds for each individual.

### 2.3. Statistical analysis

Time-to-event data analysis (Hosmer *et al*., 2008) was used to investigate the effects of DBP on daphnid survival time, age at sexual maturity (first brood), and next laying events (second brood, third, etc.). This approach enables (1) to consider events occurring during the whole study period (not only at the final time point) and (2) to take into account right-censored values, i.e. organisms for which dead/laying was not observed within the study period. Kaplan-Meier curves were used for time-to-event data representation. Log-rank tests were performed to compare survival/laying curves between the control and each condition (Harrington & Fleming, 1982). A Bonferroni correction was applied for multiple comparisons (8 or 6 comparisons, for full life cycle or adult exposure).

To investigate the effects of DBP on daphnid growth, we analysed their body size at the different measurement times. For reproduction, data were expressed as the number of neonates in the different broods (first to seventh broods) and as the total (over 25 days) number of neonates per individual-day. Indeed, this individual-day unit proved to be a suitable approach for utilising the available data and preventing bias in the estimation of ECx values (Delignette-Muller *et al*., 2014). Then, non-parametric tests were used, since the hypotheses of normality and homoscedasticity have not been validated. Wilcoxon’s tests were performed to compare each treatment to the control. A Bonferroni correction was applied for multiple comparisons (8 or 6 comparisons, for full life cycle or adult exposure).

Dose-response modelling was applied on data observed at the final time (day 25). Fitting methods were adapted to the distribution of each type of data, negative binomial for the reproduction rate per individual-day (Delignette-Muller *et al*., 2014) and normal for body length data. Effective Concentrations (EC) were calculated at the levels of 5, 10, 25 and 50% (EC_5_, EC_10_, EC_20_, EC_50_, respectively).

All the statistical analyses, tests and all graphs were performed with R 4.3.2 software (R Development Core Team, 2023), using in particular the packages *survival* 3.7-0 (Therneau *et al*., 2024) for time-to-event data analysis, *morse* 3.3.4 (Baudrot *et al*., 2024) and *drc* 3.0-1 (Ritz *et al*., 2015) for dose-response modelling and *ggplot2* 3.5.1 (Wickham *et al*., 2024) for graphics.

### 2.4. Population dynamics modelling

Results measured at the individual level were projected to the population level using an age-classified matrix population model (Leslie, 1948) with a time step of one day. The model was based on a life cycle graph of *D. magna* (Figure 2) represented with 25 one day age-classes, and assumed birth-pulses and prebreeding census (Caswell, 2001). The corresponding projection matrix ***A*** can be drawn:

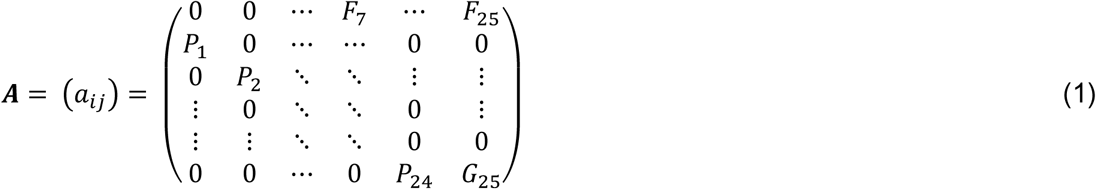

**Figure 2.**
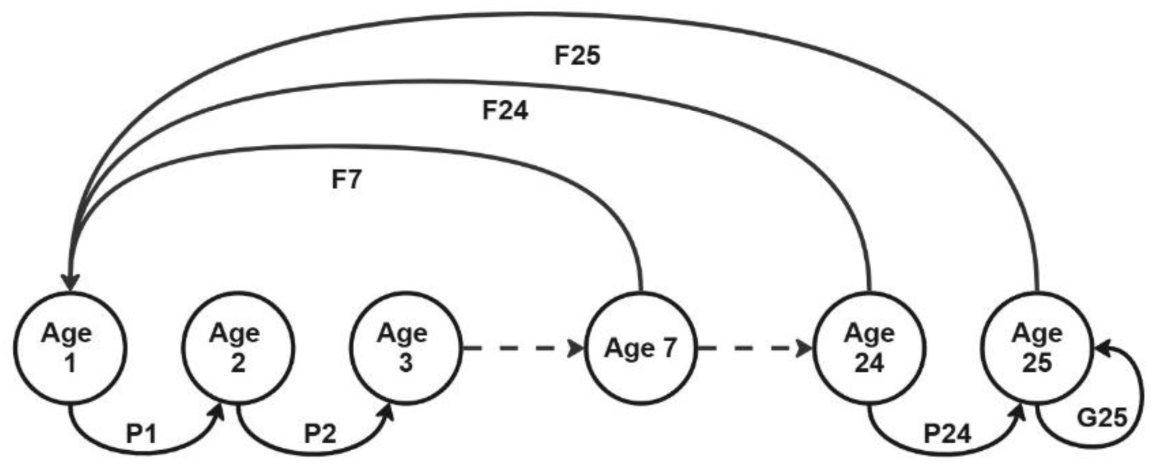
Life cycle graph of *D. magna* based on age classes representing the demographic parameters. A pre-breeding census representation in which the youngest census class is Age 1 and the fecundity corresponds to the number of neonates at time per mother multiplied by the survival of the newborns.

In this model, individuals pass from one class *i* to the next according to their condition survival rates, calculated as:

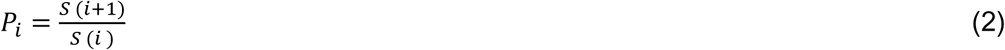

Where *S(t)* is the survivorship function as estimated from Kaplan-Meier curves. A term *G_25_*=0.95\**P_24_* was added to loop into the last age-class, as calibrated for *D. magna* (Billoir *et al*., 2007).

In the case of a prebreeding census, fecundity in age class *i* corresponds to the reproductive output of an individual upon reaching age *i* (*m_i_*) multiplied by the survival of the newborns *S*(1) (Caswell, 2001; Kendall *et al*., 2019).

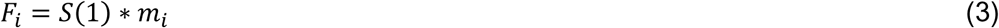

Survival and fecundity rates were specific to exposure concentration and regime (full life cycle or adult exposure). For each condition, the asymptotic Population Growth Rate (PGR) was calculated as the dominant eigenvalue (λ) of the matrix population model ***A***, and the stable age distribution as the right eigenvector ***w*** associated to λ (Caswell, 2001). PGR values provide information regarding population dynamics, since it corresponds to the multiplication factor of the population size at each time step. The stable age distribution indicates the structure of the population: scaled so that it sums to 1, it can be interpreted as the proportions of each age class in the modelled population.

We also performed elasticity analyses, in order to measure the relative sensitivity of the PGR to the different age-specific survival rates (*P_i_*) and fertility rates (*F_i_*). The terms corresponding to elasticities of λ to the entries *P_i_* and *F_i_* of matrix ***A*** can be calculated as (Caswell, 2001):

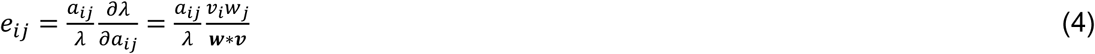

where *v_i_* and *w_j_* are the *i*th and *j*th coordinates of the left and right eigenvectors (***v*** and ***w***) of ***A***.

To assess the uncertainty associated to population modelling outputs, we used a bootstrap approach. Random re-sampling among experimental replicates (11 individuals for each condition), with replacement, was applied a thousand times. Each time, the projection matrix and the derived PGR were recalculated, resulting in Inter-Quartile Range, i.e. 25th and 75th percentiles of 1,000 PGR values.

## 3. Results

### 3.1. Effects on life history trait

#### 3.1.1. Effects of DBP on survival rate

The survival of *D. magna* exposed to different DBP concentrations is represented as Kaplan-Meier curves in Figure S1 (supplementary data) and Figure 3 for adult and full life cycle exposure regimes, respectively. Survival proportions at some target times (days 0, 5, 10, 15, 20 and 25) and their associated uncertainty are also shown in Figure S2. For both exposure regimes, no mortality was observed in the control condition at the end of the experiment.

**Figure 3.**
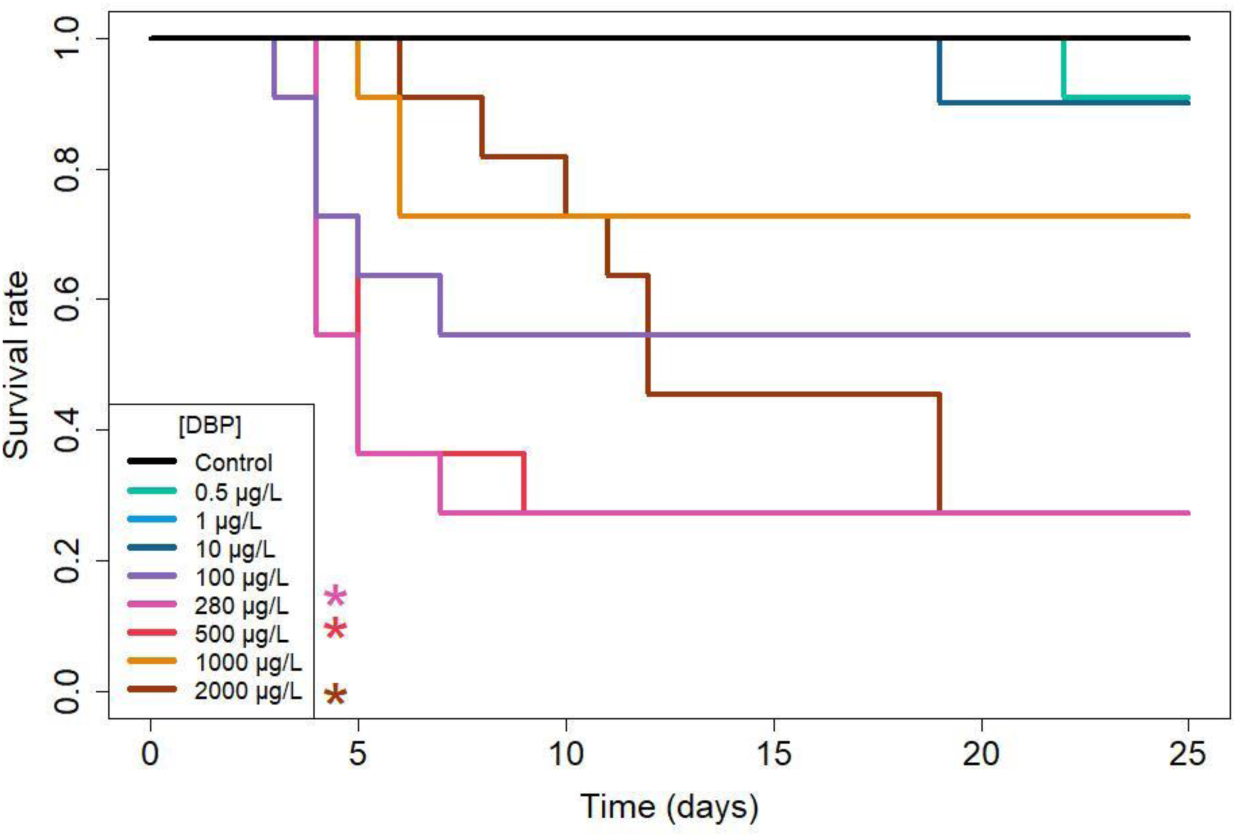
Survival rate of *D. magna* exposed to DBP under full life cycle regime (n=11 for each concentration). *p ≤ 0.05 according to log-rank tests comparing each treatment to the control condition (p-value adjusted by a Bonferroni correction).

No effect of DBP on survival was reported in the adult exposure regime (log-rank test; p > 0.05; Figure S1). The maximum mortality was limited to 18% on day 25 in the 10 µg/L treatment.

In contrast, a significant effect of DBP was reported on the full life cycle exposure regime. While the exposure to the three lowest concentrations (0.5, 1, 10 µg/L) resulted in less than 10% mortality at the end of the experiment, daphnids exposed to 280, 500 and 2000 µg/L showed significantly higher mortality compared to the control (72% at day 25; log rank tests; p > 0.05; Figure 3). Mortality displayed a non-monotonous dose-response relationship, with higher mortality at 280 and 500 µg/L (72% at the end of the experiment) than at 1000 µg/L (27%). Similar mortality dynamics were observed for exposure to 100, 280, 500 and 1000 µg/L with juvenile mortality (between days 4 and 7), followed by no mortality from day 9 onwards. Notably, exposure to 2000 µg/L resulted in a gradual increase in daphnid mortality from day 6 to day 19 of the experiment, reaching a mortality rate of 72%.

#### 3.1.2. Effects of DBP on size

The effect of DBP on daphnid size is shown in Figure S3 for the adult exposure regime (size at days 0, 7, 13 and 25) and in Figure 4 for the full life cycle exposure regime (size at days 0, 2, 4, 7, 13 and 25). For both exposure regimes, under the control condition, daphnids grew from 0.82 - 0.89 mm (day 0) to 3.7 - 4 mm (day 25).

**Figure 4.**
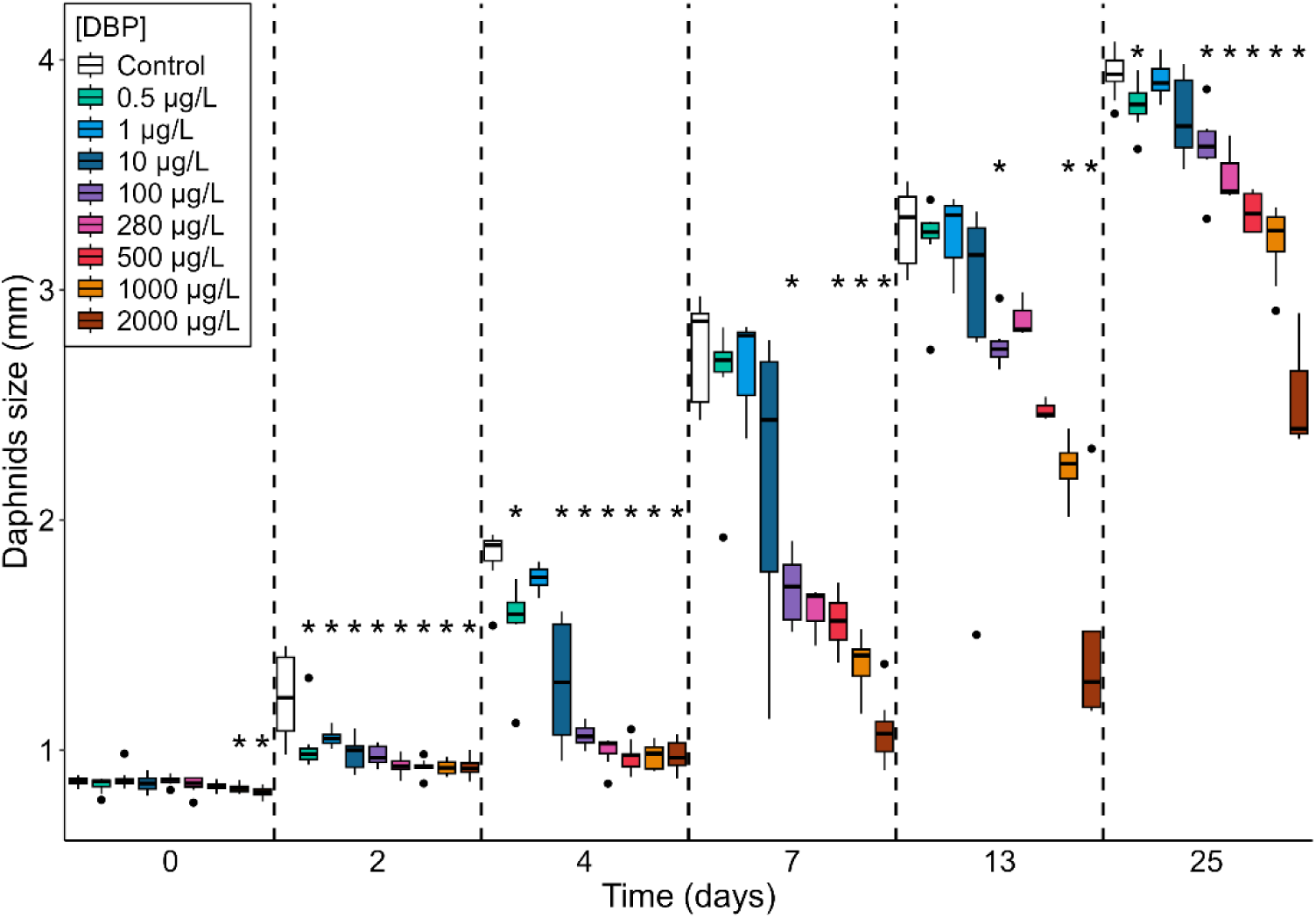
Size of *D. magna* exposed to DBP under full life cycle regime and measured at day 0, 2, 4, 7, 13 and 25 on all surviving individuals. *p ≤ 0.05 according to Wilcoxon tests comparing each treatment to the control condition, n = 11 (p-value adjusted by a Bonferroni correction). The horizontal line represents the median, the boxes range from the 25th to the 75th percentile and the whiskers extend to the last values within 1.5 times the inter-quartile range with black dots representing outliers.

No effect of DBP was reported on daphnid size in the adult exposure regime, as daphnids exhibited similar sizes across all DBP concentrations (no effects of DBP on size at day 0, 7, 13 and 25; Wilcoxon test; p > 0.05; Figure S3).

The full life cycle DBP exposure regime induced a significant effect on daphnids growth. Compared to control, daphnid size was significantly reduced by 3.5% to 5.3% for the two highest concentration, on day 0 (Wilcoxon test; p < 0.05; Figure 4). On day 2, size was significantly reduced by 19 to 25% from 0.5 to 2000 µg/L (test de Wilcoxon; p < 0,05; Figure 4). On day 4, 0.5 µg/L was the lowest concentration to cause a significant size reduction (by 15%), with significant reductions ranging from 29 to 47 % from 10 to 2000 µg/L. This dose-dependent size reduction attenuated over time, with significant size reductions of 37 to 60 % from 100 to 2000 µg/L on day 7, and by 15 to 54 % from 100 to 2000 µg/L on day 13 (Wilcoxon test; p < 0.05; Figure 4). By the end of the experiment, on day 25, 0.5 µg/L induced a size reduction of 3.4%, and the concentrations from 100 to 2000 µg/L lead to a significant size reduction, compared to control (Wilcoxon test; p < 0.05; Figure 4).

EC_5, 10, 20_ and _50_ estimated for daphnid size at day 4, 7 and 25 under the full life cycle exposure regime are provided in Table 1. The EC at day 4 are very low (for example EC_10_-4d: 0.25 µg/L), while at day 7 the sensitivity decreases (EC_10_-7d: 1.16 µg/L). Finally, at day 25 the EC_10_-25d is 177.76 µg/L illustrating a progressive decrease of the sensitivity along the time of exposure.

**Table 1.** Summary table showing the EC_5, 10, 20_ and _50_ values evaluated for growth of *D. magna* exposed to DBP in full life cycle regime on the days 4, 7 and 25, with 95% confidence intervals.

|  | EC<sub>5</sub> | EC<sub>10</sub> | EC<sub>20</sub> | EC<sub>50</sub> |
| --- | --- | --- | --- | --- |
| **Day 4** | 0.08<br>[0 - 0.23] | 0.25<br>[0 - 0.59] | 0.82<br>[0.03 - 1.61] | 6.22<br>[2.34 - 10.11] |
| **Day 7** | 0.33<br>[0 - 0.90] | 1.16<br>[0 - 2.77] | 4.46<br>[0 - 9.31] | 44.85<br>[14.19 - 75.51] |
| **Day 25** | 37.62<br>[0 - 91.80] | 79.22<br>[0 - 167.49] | 177.76<br>[41.24 - 314.28] | 707.79<br>[502.36 - 913.22] |

#### 3.1.3. Effects of DBP on reproduction

The effect of DBP on daphnid reproduction (sexual maturity and total number of neonates per individual-day) is shown in Figure S4 for the adult exposure regime and in Figure 5 for full life cycle exposure regime. In control conditions, the first brood of daphnids occurred between days 5 and 8 in both exposure regimes. It is important to note that in the adult exposure regime, daphnids were exposed to DBP starting on day 14, which was later in their development relative to their first brood. The timing of the first brood was consistent across all treatments (log-rank test; p > 0.05; Figure S4.A).

**Figure 5.**
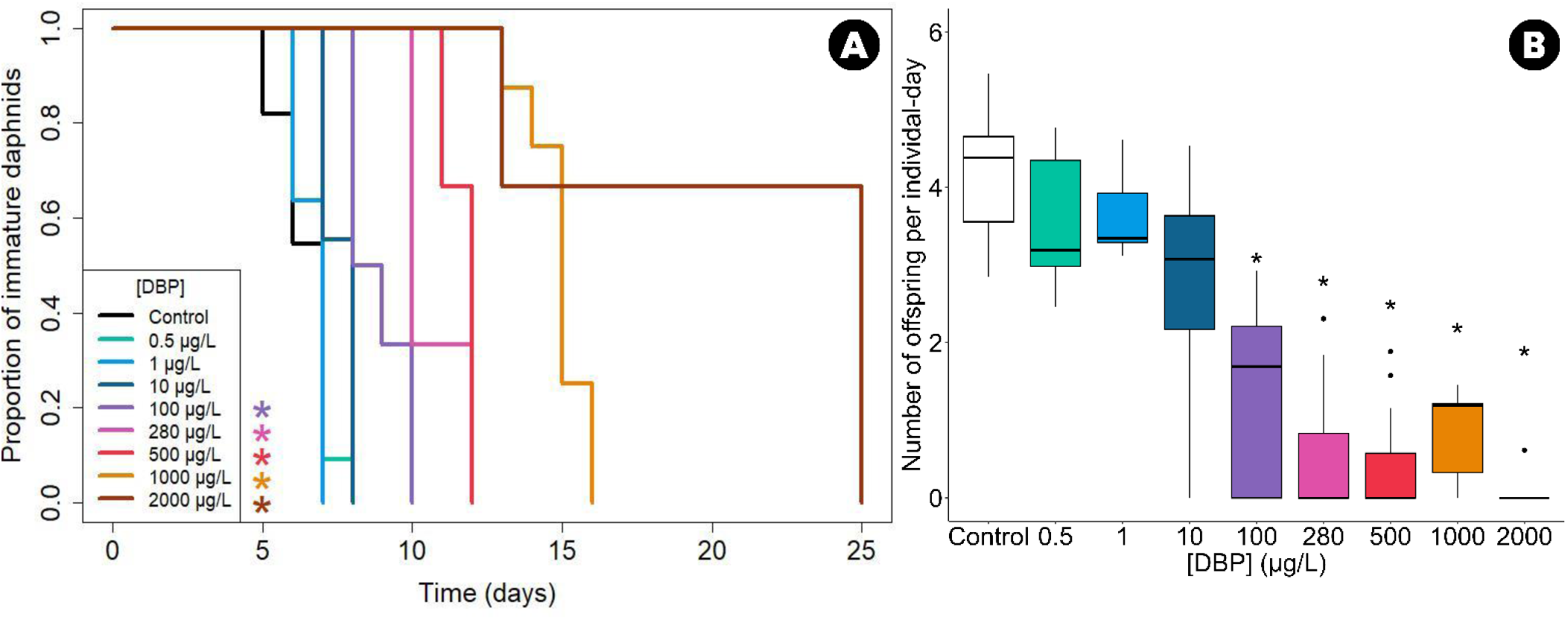
Reproduction of daphnids exposed to DBP under full life cycle regime. (A) Time to sexual maturity of *D. magna* exposed to DBP, expressed as the proportion of daphnids that have not yet laid their first brood among surviving individuals (n=11 for each concentration). *p ≤ 0.05 according to log-rank tests comparing each treatment to the control condition (p-value adjusted by a Bonferroni correction). (B) Fecundity of *D. magna* exposed to DBP, expressed as number of offspring per individual-day as a function of concentration (n=11 for each concentration). *p ≤ 0.05 according to Wilcoxon tests comparing each treatment to the control condition (p adjusted by a Bonferroni correction). The horizontal line represents the median, the boxes range from the 25th to the 75th percentile and the whiskers extend to the last values within 1.5 times the inter-quartile range with black dots representing outliers.

In the full life cycle DBP exposure regime, the timing of the first brood was significantly delayed, following a monotonous concentration-dependent relationship: higher exposure concentrations resulted in greater delays (Figure 5.A). Daphnids exposed to the three lowest tested concentrations (0.5, 1 and 10 µg/L) showed no significant effect compared to the control (first brood between day 6 and 8, log-rank test; p > 0.05). However, those exposed to concentrations higher than 100 µg/L showed significant delays in the timing of the first brood (log-rank tests; p < 0.05). At 100, 280, 500, 1000 and 2000 µg/L, the first brood was observed on days 8-10, 10-12, 11-12, 13-16 and 13-25, respectively. Subsequent broods were also delayed (Figures S8), with no effect on the intervals between broods.

On average, 102 neonates per daphnid were produced over 25 days in the controls, corresponding to 4.3 neonates per individual-day. Under the adult exposure regime, DBP had no significant effect on the number of neonates per individual-day (Wilcoxon tests; p > 0.05; Figure S4.B). However, the full life cycle exposure regime significantly reduced the number of neonates produced per individual-day, following a monotonous concentration-dependent relationship (Figure 5.B). Daphnids exposed to the three lowest concentrations (0.5, 1 and 10 µg/L) showed a similar number of neonates per individual-day compared to the control (log-rank test; p > 0.05). In contrast, those exposed to concentrations higher than 100 µg/L showed a significant reduction in neonate production per individual-day (log-rank test; p<0.05); by 45%, 53%, 62%, 72% and 85% under exposure to 100, 280, 500, 1000 and 2000 µg/L, respectively. Under the full life cycle exposure regime, for reproduction rate per individual-day, we assessed an EC_5_-25d of 0.35 [0.04 - 2.30] µg/L, an EC_10_-25d of 1.01 [0.16 - 5.04] µg/L, an EC_20_-25d of 3.14 [0.72 - 11.83] µg/L and an EC_50_-25d of 22.16 [8.49 - 53.35] µg/L (95% confidence intervals).

### 3.2. Impact of DBP on population dynamics

The results obtained through population modelling illustrated how combined effects on several individual endpoints may propagate to the population level.

As a population endpoint, the population growth rate (PGR) is shown in Figure 6.A for both exposure regimes. The value of the PGR in the control condition was 1.41 (IQR 1.4-1.42) per day, representing a population growth of 41% per day, and meaning that our control population had a very high growth potential. Results derived from the adult exposure regime data were similar to the control condition for all exposure concentrations, with PGR values between 1.39 and 1.41. Overlapping IQRs indicated that the PGR was not different between exposed and control conditions. Results derived from the full life cycle exposure regime data highlighted a progressive decrease in the PGR with increasing exposure concentrations. Population-level impacts were limited at the lowest tested concentrations. Exposure to 100 µg/L resulted in a reduction in the PGR of 12.4% compared to the control. The PGR was even lower for higher concentrations, such as 500 µg/L and 2000 µg/L, with reduction of 19.7% and 27.7%, respectively.

**Figure 6.**
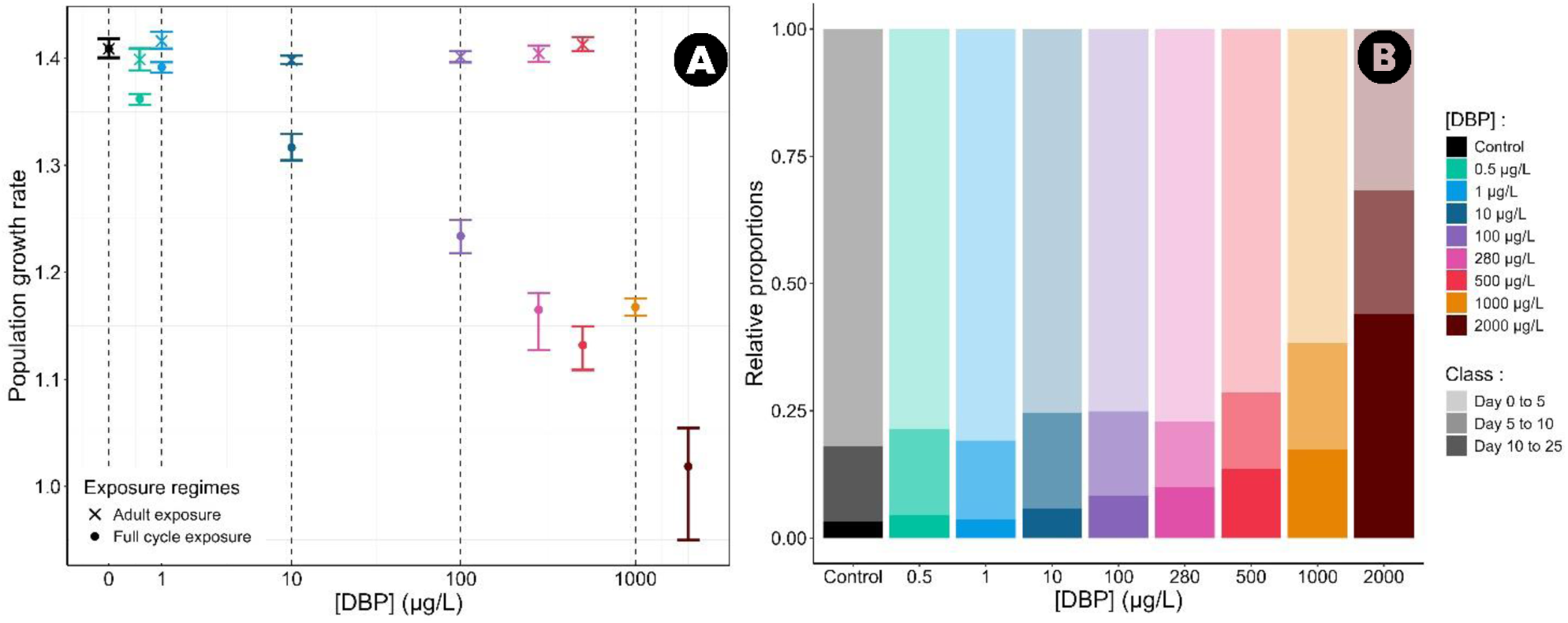
(A) Population Growth Rate (PGR) of *D. magna* laboratory population for each DBP exposure concentration under both exposure regimes. The points represent the median PGR values, while the upper and lower bounds correspond to the 25th and 75th percentiles of the 1000 PGR values obtained through bootstrapping. (B) Stable age distribution of *D. magna* laboratory populations for each DBP concentration under full life cycle exposure. The stable age distribution is represented in three age classes (day 0 to 5, day 5 to 10 and day 10 to 25 days; faded colors).

We investigated how DBP exposure may affect the stable age distribution, which represents the long-term structure of the modelled population. The results were cut into three age classes (0 to 5 days old, 5 to 10 days old and 10 to 25 days old) (Figure 6.B, Figure S5) to represent the proportions of juveniles, young adults, and adults, respectively. In the control, the population was predominantly made up of juveniles (82%), with only 15% of young adults and 3% of adults. The population structures for each exposure condition under adult exposure regime were similar to those of the control condition, with very small changes in the stable age distribution (less than 1%) compared to the control values (Figure S5). Under the full life cycle exposure regime, as the concentration increased, the stable age distribution was progressively impacted, with an increase in the adult proportion and a decrease in the juvenile proportion (Figure 6.B). At low concentrations, the stable age distribution was similar to the control, with a predominant juvenile proportion (75%). However, as the exposure concentration increased, the proportion of adults also increased.

Elasticity analysis of our model represented the relative sensitivity of the PGR to changes in life history traits (namely age-specific survival rates and fecundity rates). The results are shown in Figure S6 for both exposure regimes. For the modelled population in the control condition, the PGR was overall equally sensitive to survival and fecundity rates, with sums of elasticity coefficients at 0.504 and 0.496, respectively.

The survival of the first age class had the highest elasticity coefficient (0.25), with this value rapidly decreasing in subsequent age classes. Similarly, the fecundity rates of young mature daphnids had the highest elasticity coefficients (0.11 for the eighth age class), whereas the PGR was less sensitive to the fecundity rates of older classes. For the adult exposure regime, we observed similar patterns to those of the control condition, regardless of the exposure concentration. For the full life cycle exposure regime, except for 2000 µg/L, we observed similar patterns for survival rates with a lower contribution to the survival rate for the first few days and a strong contribution to the survival rate on the day 25 (0.4). Again, the PGR was especially sensitive to the fecundity rates of the first mature age classes. However, as a delay in the onset of the reproduction was observed in daphnids under the full life cycle DBP exposure regime, the elasticity coefficients of fecundity rates became positive at later age classes as the exposure concentration increased: for 10, 100, 280, 500, 1000 and 2000 µg/L, the first day with a positive fecundity rate was 10, 11, 13, 14, 16 and 16, respectively.

## 4. Discussion

The effects of a wide range of DBP exposures concentration on *D. magna*’s life history traits (survival, growth, reproduction) were assessed following two exposure regimes: (1) from the beginning of embryonic development (full life cycle exposure regime) or (2) from the adult stage from the third brood (adult exposure regime). In order to assess long-term consequences, population-level effects were estimated on various demographic metrics using a matrix modelling approach.

### 4.1. Effects on daphnid survival

In the present study, the adult exposure regime from 0.5 to 500 µg/L of DBP exerted no chronic effect on daphnid survival whereas survival of organisms exposed to the same DBPs concentrations according to the full life cycle exposure regime was significantly reduced (significant for exposure to 280, 500 and 2000 µg/L of DBP). We observed that DBP had an early effect on survival at concentrations starting from 100 to 1000 µg/L, with mortality occurring in daphnids aged 4 to 7 days (Figure 3). The non-monotonic dose-response relationship we observed for survival, with higher mortality rates at intermediate concentrations (280 and 500 μg/L) than at 1000 μg/L, constitutes an intriguing result. This observation can potentially be attributed to uncertainty in estimating survival rate based on a limited sample size (Figure S8). However, while these effects have never been documented for DBP on *D. magna*, similar patterns have been reported in other studies. An exposure of daphnia neonate to *Bacillus thuringiensis* resulted in a non-monotonic dose-response for a 24-hour exposure (de Souza Machado *et al*., 2017). Similarly, McMahon *et al*. (2011) reported non-monotonic dose-responses on survival for three amphibian species exposed to chlorothalonil, with low and high levels causing significantly higher mortality than intermediate levels and controls. Mechanistically, this non-monotonic dose-response relationship may result from the combined action of two or more modes of action, with molecular mechanisms differentially affecting the response according to concentration. At intermediate concentrations, DBP may act primarily through specific pathways affecting juvenile development and survival, while at higher concentrations, additional compensatory mechanisms could reduce mortality effect. It should be noted that assessing lethal effects was not the primary objective of this study. Survival was monitored as part of the life-history traits evaluation (individual-day observations) and to verify the health status of control organisms. Investigating lethal effects would require an experimental design with a substantially higher number of replicates.

The acute toxicity of DBP in *D. magna* has been described in several studies, with EC_50_-24h values (Effective Concentration values for 50% of organisms after 24 h exposure) ranging from 3.04 to 10.35 mg/L, and EC_50_-48h values ranging from 2.55 and 5.2 mg/L (McCarthy & Whitmore, 1985b; Adams *et al*., 1995b; G. Huang *et al*., 1998; Jonsson & Baun, 2003; B. Huang *et al*., 2016b; Wei *et al*., 2018b; Seyoum & Pradhan, 2019b; Shen *et al*., 2019b). To our knowledge, the chronic toxicity of DBP on *D. magna*’s survival has been investigated in only six studies, with exposure durations ranging from 16 to 60 days, and without embryonic exposure (exposure starting with neonates <24h; McCarthy & Whitmore, 1985b; Defoe *et al*., 1990b; Rhodes *et al*., 1995; Wei *et al*., 2018b; Seyoum & Pradhan, 2019; Jin *et al*., 2024). McCarthy & Whitmore (1985) observed 30% mortality after a 16-day exposure to 1.8 mg/L, Defoe *et al*. (1990) reported an LC50-21d of 1.91 mg/L, and Rhodes *et al*. (1995) reported 69% mortality after a 21-day exposure to 2.5 mg/L. Similarly, Seyoum & Pradhan (2019) observed an effect of 2.78 mg/L on daphnid survival, but only after 30 days of exposure, resulting in a reduction in the average lifespan of exposed daphnids, while they were unaffected by exposure to 278 µg/L. Based on the limited literature on the effects of DBP on *D. magna*, we noticed that: (1) the effect of DBP on survival appears at concentrations in the mg/L range, which is far above environmental concentrations (see introduction), and (2) the effects seem to depend on exposure conditions. Thus, our results are the first to highlight chronic effects of DBP on *D. magna*’s survival below the mg/L range, with the noteworthy particularity that exposure started at the beginning of embryonic development (full life cycle exposure regime). The influence of the exposure window on endpoint response has been reported in many publications, showing that the effect is more severe when the embryonic stage is exposed (Massarin *et al*., 2010; Plaire *et al*., 2013). Similarly, Shen *et al*. (2019) reported that the exposure window influences DBP EC_50_-48h in *D. magna*, with neonates exhibiting higher sensitivity compared to adults (2.83 *vs* 4.92 mg/L, respectively).

### 4.2. Effects on daphnid growth

Similarly to survival, growth was not affected by DBP in daphnids subjected to the adult exposure regime. This observation makes sense, as the exposure had begun after day 14, when most energy investment in growth is already completed.

In contrast, under the full life cycle exposure regime, daphnid sizes were significantly reduced from 1000 µg/L at day 0 (juvenile release day), showing that DBP affected growth during embryonic development (Figure 4). While no specific data appears to be available on the effects of DBP on daphnid embryonic development, Li *et al*. (2021) reported that another phthalate, butyl benzyl phthalate (BBP), significantly impacted daphnid eye development, resulting in abnormal compound eyes in first instar juveniles exposed to 600 and 1200 µg BBP/L. In other species, DBP has been shown to reduce development rate in the marine shrimp (*Palaemonetes pugio*) (Laughlin *et al*., 1978: at 100 and 500 µg/L) and to reduce body size in the zebrafish (*Danio rerio*) after embryonic exposure (Sun & Li, 2019: by 5-10% for exposition to 500 and 1000 µg/L).

On days 2 and 4, under full life cycle exposure regime, we observed significant reductions in daphnid size starting at 0.5 µg/L. These concentration-dependent size reductions persisted over time (days 7 to 25) but gradually decreased at lower concentrations (Figure 4). The few studies focusing on the effect of DBP on daphnids growth in neonates (<24h) mainly reported an absence of effect on daphnid size after 21 days of DBP exposure (Huang *et al*., 1999: from 500 to 4000 µg/L; Wei *et al*., 2018: from 70 to 480 µg/L). However, Seyoum & Pradhan (2019) reported significantly smaller sizes after 14 days of exposure to 2.78 mg/L, and Wei *et al*. (2018) observed that embryonic exposure to 70 µg/L DBP in daphnids, followed by transfer to clean water after hatching, significantly reduced growth rates. Plaire *et al*. (2013) also showed that the effects of uranium on the life cycle of *D. magna* were more pronounced in organisms exposed at the embryonic stage compared to those after hatching. Thus, compared to the literature, the greater effect of DBP on daphnid growth we reported could be mainly attributed to embryonic exposure.

According to our data, DBP seems to affect daphnid growth during the early stages of development, with these effects diminishing over time at the lowest concentrations. The 10 µg/L concentration appeared to be pivotal, showing high size variability across all measured times (Figure 4), suggesting differential responses between sensitive and resistant individuals. An increase in variability for a parameter may reflect the organisms’ ability to acclimate or an early response to stress and should be more frequently considered in studies (Devin *et al*., 2014).

### 4.3. Effects on daphnid reproduction

As with survival and growth, all reproductive endpoints were unaffected by DBP when daphnids were subjected to the adult exposure regime. Under the full life cycle exposure regime, we observed a significantly delayed age of first brood and subsequent broods starting from 100 µg/L, resulting in a reduced number of broods over the 25-day period (Figure S7). There were significant reductions in the number of neonates produced per individual-day, starting at 100 µg/L, in a concentration-effect monotonic relationship along the exposure gradient (Figure 5). Additionally, when looking at each brood individually, we observed a trend of reduced numbers of neonates with increasing DBP concentrations (Figure S8).

In the literature, the effects on chronic DBP exposure on daphnid reproduction are quite variable. Huang *et al*. (1999) observed a delay of 2 to 3 days in the first brood at DBP exposure of 4000 µg/L, and Seyoum & Pradhan. (2019) showed a delay of one day at 278 and 2780 µg/L. Authors have highlighted either 1) a reduction in the total number of neonates per daphnid (Defoe *et al*., 1990: IC_50_ 21-day of 1640 µg/L, McCarthy & Whitmore, 1985: 1800 and 3200 µg/L) 2) no effect (Rhodes *et al*., 1995: 70 to 2500 µg/L) or 3) an increase in the total number of neonates per daphnid (Wei *et al*., 2018: 70 to 480 µg/L), or per brood (Wei *et al*., 2018: 270 & 420 µg/L). Authors exposing daphnids only during the embryonic stage reported no effects of DBP (Wei *et al*., 2018: 70 to 480 µg/L). Compared to this literature, as for growth, the noteworthy decrease in reproduction we reported could be attributed to embryonic exposure.

Our adult exposure regime aimed to investigate direct effects of DBP exposure on reproductive endpoints, as investment in growth is highly reduced at this stage. Our results instead suggested that the effects on reproduction originated before the reproductive period, either as a consequence of delayed growth (influencing age of maturity) and/or impairment in reproductive organ development.

### 4.4. Endpoint dependency and sensitivity

Under optimal developmental conditions (temperature, food), *Daphnia* reaches sexual maturity between days 5 and 10, after molting between 4 and 5 times (Ebert, 2005). The molting process paces the growth and reproduction of *D. magna*. In the present study, the effect of DBP on the number of offspring per individual-day seems to be mainly due to the delay in first brood. Since this delay is very marked while the interval between two brood events remains similar regardless of the brood event and exposure concentration tested (Figure S7), the delay in the first brood may be associated with a lack of growth in the daphnids, likely due to a lack of molting (e.g., daphnids exposed to 500 µg/L for 13 days are similar in size to control daphnids at day 7). Furthermore, there is a positive correlation between daphnid size and the number of juveniles produced (Boersma, 1997). Shifting the date of first brood thus reduces the number of brood events achievable over the duration of the experiment (25 days) and *de facto*, the number of juveniles produced during the experiment per individual-day. Moreover, our results showed a declining trend in the number of juveniles produced per brood and per individual with increasing DBP exposure concentration (Figure S8). Thus, at the end of the experiment, the number of juveniles per individual-day should be most sensitive trait, given the attenuation of growth effects over time and the accumulation of reproductive effects. Over a long exposure period, the total number of juveniles produced by daphnids is influenced by several factors: (1) the growth of daphnids, which is a prerequisite for reaching sexual maturity and determines the total number of broods during the experiment, (2) the number of juveniles produced in each brood, which can amplify the total juvenile production, and (3) the size of the females, with larger females producing more juveniles. In the present paper, when considering the responses at the end of the experiment (25 days), reproduction (number of juveniles produced per individual-day) is the most sensitive endpoint (EC_10_-25d of 1.01 µg/L for number of juveniles per individual days vs 79.22 µg/L for growth). In daphnid chronic bioassays, reproduction is often reported as the most sensitive endpoint (Mark & Solbé, 1998; Toumi *et al*., 2015). However, considering all exposure times and endpoints, while reproduction events has not yet started, growth endpoint at day 4 is the most sensitive endpoint (EC_10_-4d of 0.25 µg/L) followed by growth endpoint at day 7 (EC_10_-7d of 1.16 µg/L; similar to EC_10_-25d on number of juveniles produced per individual-day), suggesting the main and strong early impacts of DBP on growth. Our results confirm the strong relationship between growth impairment and reproductive effects, as highlighted by previous studies on *D. magna* (Green, 1956; Barata *et al*., 2001). Even if reproduction is a conclusive endpoint for assessing sensitive effects in the case of chronic exposure, our results show that it is important to also consider the size of organisms at different times of exposure to identify the sensitivity and early responses of organisms to contaminants.

### 4.5. Population impacts

The value of the population growth rate (PGR) in the control condition was estimated at 1.41 (IQR 1.4-1.42) per day, meaning that our control population had a very high growth potential. This value is consistent with other studies on laboratory *D. magna*, like Massarin (2010) who has reported a PGR of 1.35 per day, Pieters *et al*. (2006) and Heugens *et al*. (2006) who have reported instantaneous per capita growth rates (noted *r* and equal to log (PGR); Stevens, 2009) corresponding to PGR of 1.35 and 1.52 per day, respectively.

Population modelling aimed to assess the combined effects of DBP exposure on lethal and sublethal endpoints. As for life history traits, population endpoints were unaffected by DBP when daphnids were exposed only as adults. However, under the full life cycle regime, we observed a progressive decrease in the PGR with increasing exposure concentration, with a 19.7% reduction in the PGR at 500 µg/L compared to the control (Figure 6). For example, over three days (one reproductive cycle), the theoretical control population would multiply by 2.8 (1.41^3^), whereas the population exposed to 500 µg/L would only multiply by 1.45 (1.13^3^). Nevertheless, the PGR appeared less sensitive than the individual endpoints (EC_20-7d_ growth: 4.46 µg/L; EC_20-25d_ reproduction: 3.14 µg/L). According to elasticity analyses, the PGR was mostly influenced by the survival rates of the first few age classes and the fecundity rates of the first few age classes able to reproduce. Previous studies on *D. magna* have highlighted that PGR is especially driven by juvenile and maturity-related parameters (Billoir *et al*., 2007; Alonzo *et al*., 2016). Strikingly, the effects of DBP were quite strong at these stages.

In ecological risk assessment, the population level is acknowledged to be more relevant than the organismal level (Forbes *et al*., 2011). In this context, ecological risk assessment would strongly improve with the consideration of realistic environmental conditions. Standardised conditions, as used in our study, do not represent the natural situation where population responses to contaminants may vary under limited food levels or suboptimal temperatures (Heugens *et al*., 2006). Density-dependence has also been shown to influence *D. magna* population dynamics (Preuss *et al*., 2009). However, in the present study, the matrix model is used not as a tool for forecasting population dynamics but as a projection tool at the population level (Caswell, 2001). This approach allows for an integrated analysis of the responses to DBP observed through life history traits and the relative sensitivity of PGR to survival and fecundity rates. Another important point to consider in populational approaches, is the potential for multi/transgenerational effects, which our study did not address. Investigating disruptions across multiple generations would enable to project more precisely the long-term effect of contaminants on populations (Biron *et al*., 2012; Wu *et al*., 2025).

### 4.6. Sources of variability

Given the variability in DBP effects reported in the literature and this study, it seems essential to carefully examine the conditions, methods and biological materials used in experiments from a risk assessment perspective. Differences in responses to DBP may be due to inherent differences between daphnid populations or strains, arising from genotype differences (Barata & Baird, 1998; Picado *et al*., 2007; Toumi *et al*., 2013), exposure regimes (Massarin *et al*., 2010; Plaire *et al*., 2013) or experimental conditions that influence speciation and bioavailability (e.g. resource quantity: Baird & Barata, 1998; resource quality: Ruiz *et al*., 2022). For example, it is noteworthy that control *D. magna* size is highly variable between studies (at day 7: close to 0.8 mm in Seyoum & Pradhan (2019), 2.3 mm in this study). With such size differences, individuals will likely respond differently to contaminants. Chenon *et al*. (2000) discussed the issue of genetic homogeneity in strains used by laboratories testing parthenogenetically reproducing animals. Ruiz *et al*. (2022) reported significant effects of food resource quality on *D. magna* survival, growth, reproduction, and sensitivity to stressors. Genotype and culture conditions should be specified for laboratories conducting standard *D. magna* bioassays (Baird *et al*., 1990; Toumi *et al*., 2013), although this is rarely done, even though it is mentioned in the *ad hoc* ISO standard (ISO 6341. (1996) as a way to improve inter-laboratory reproducibility (Férard & Férard, 2013). It is also important to note that exposure conditions (chemical composition of the environment, density of organisms, solvent utilization) may also contribute to differences in results and must be considered.

## Conclusion

DBP is a ubiquitous phthalate in freshwater environments. The present study is the first to assess the effects of DBP on the life history traits of *Daphnia magna* over a wide range of concentrations, without solvents, and including embryonic development. Our findings suggest that the effects of DBP may have been underestimated in the past and more importantly, that the exposure regime is a key factor in assessing the effects of a contaminant on the life history traits of *D. magna*. While no significant effects were observed in organisms exposed from the third brood onward, exposure from embryonic development (full life cycle regime) led to significant and sensitive effects on survival, growth and. Projecting the individual trajectories of daphnids exposed to a full life cycle regime reveals a progressive decrease in population growth rate, as well as population aging, across the concentration range. This integrated approach provides a better understanding of how effects on survival and fertility shape population dynamics.

## Data, scripts, code, and supplementary information availability

- All the data and scripts are available online: https://doi.org/10.57745/ZKF4CD

## Funding

This work was supported by the ANR under grant ANR-21-CE34-0003 (JCJC Chroco project).

## Acknowledgements

This work was partly done in the “Pôle de compétences en biologie environnementale” and the “Pôle de compétences en chimie analytique environnementale”, ANATELo, LIEC laboratory, UMR 7360 CNRS – Université de Lorraine.

We would thank the Organic Geochemistry laboratory of GeoRessources in Nancy, included in ANATELo and ReGEF facility, and which is partly funded by the Région Lorraine and the European Community through the FEDER program.

We would like to express our thanks to Céline Simon for experimental support and David Billet for performing DBP measurements in stock solutions, improving the quality of this research.

A preprint version of this article has been peer-reviewed and recommended by PCI Ecotox Env Chem (https://doi.org/10.24072/pci.ecotoxenvchem.100248).

## Notes

https://doi.org/10.57745/ZKF4CD

### Competing Interest Statement

The authors have declared no competing interest.

### Summary of Updates

This is the final version after peer-reviewing.

