## Supplementary data for "Effects of dibutyl phthalate (DBP) on life history traits and population dynamics of *Daphnia magna*: comparison of two exposure regimes"

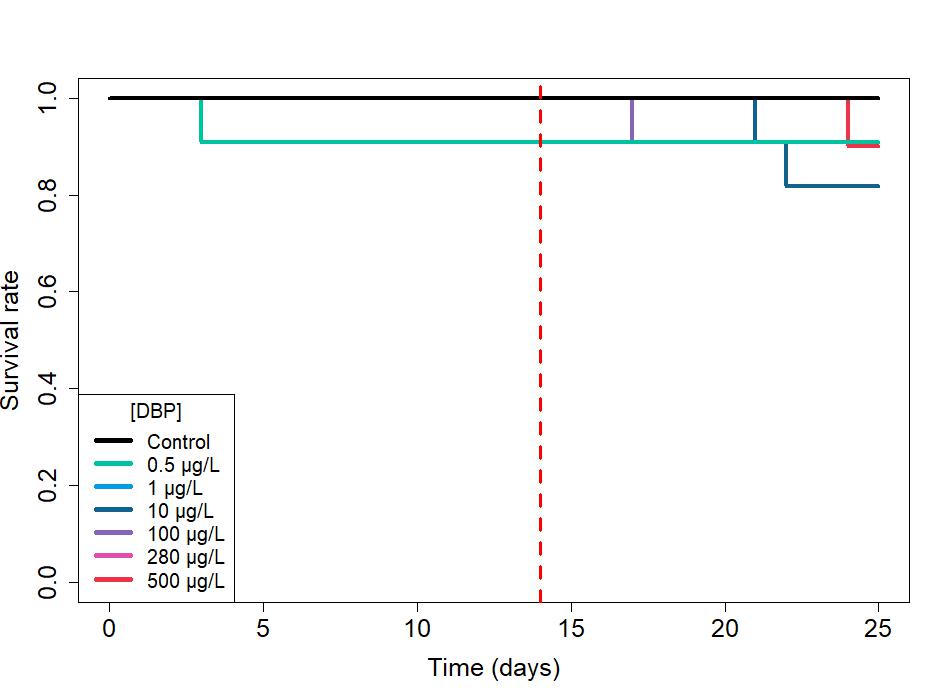
**Figures in supplementary data**

**Figure S1.** Survival rate of *D. magna* exposed to DBP under adult exposure regime (n=11 for each concentration). *p ≤ 0.05 according to log-rank tests comparing each treatment to the control condition (p-value adjusted by a Bonferroni correction). The red dotted line represents the onset of exposure to DBP.

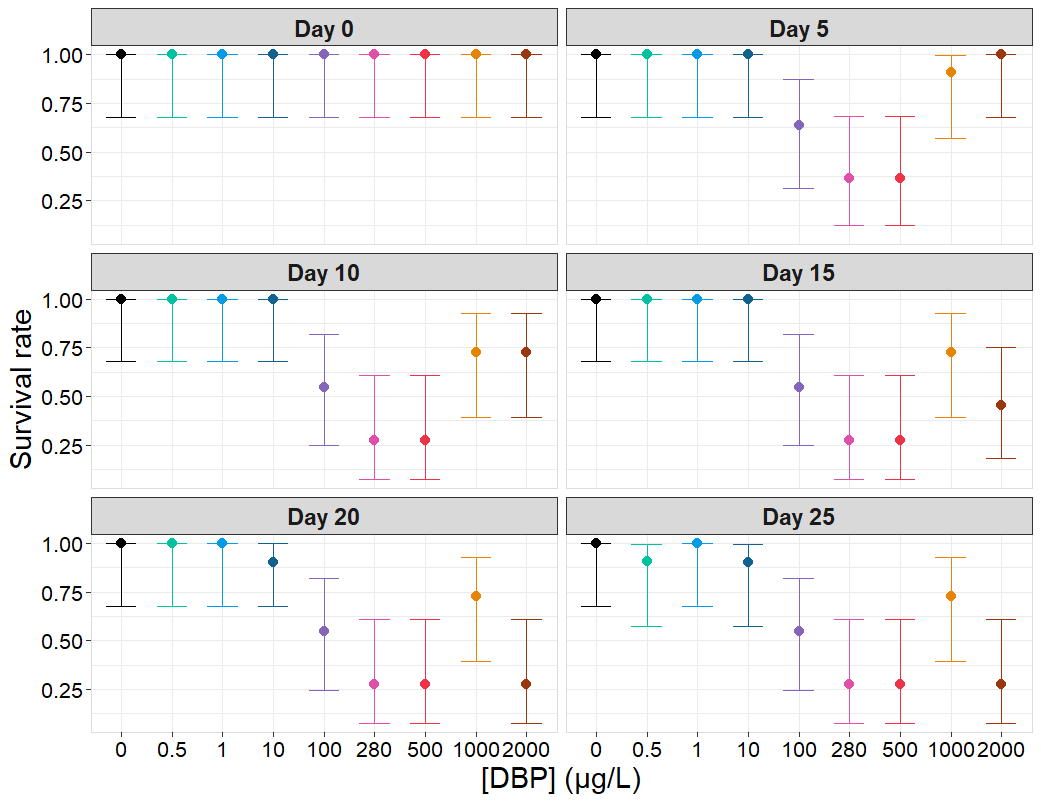

**Figure S2.** Survival rate of *D. magna* exposed to DBP under full life cycle regime at specific time points (Days 0, 5, 10, 15, 20 and 25). Error bars show the range of statistical uncertainty in survival estimates (95% confidence interval; n=11 for each concentration).

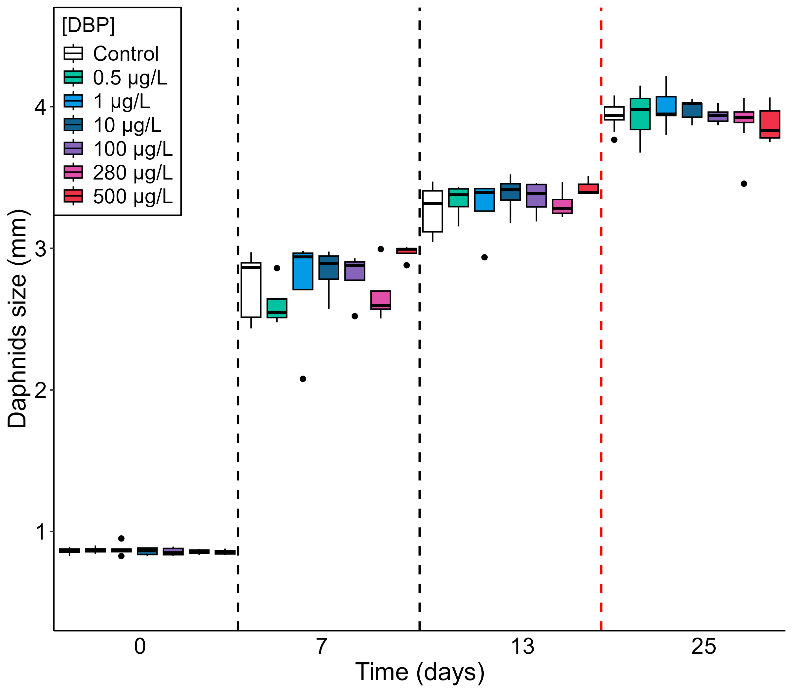

**Figure S3.** Size of *D. magna* exposed to DBP under adult regime and measured at day 0, 7, 13 and 25 on all surviving individuals. *p ≤ 0.05 according to Wilcoxon tests comparing each treatment to the control condition, n = 11 (p-value adjusted by a Bonferroni correction). The horizontal line represents the median, the boxes range from the 25th to the 75th percentile and the whiskers extend to the last values within 1.5 times the inter-quartile range with black dots representing outliers. The red dotted line represents the onset of exposure to DBP.

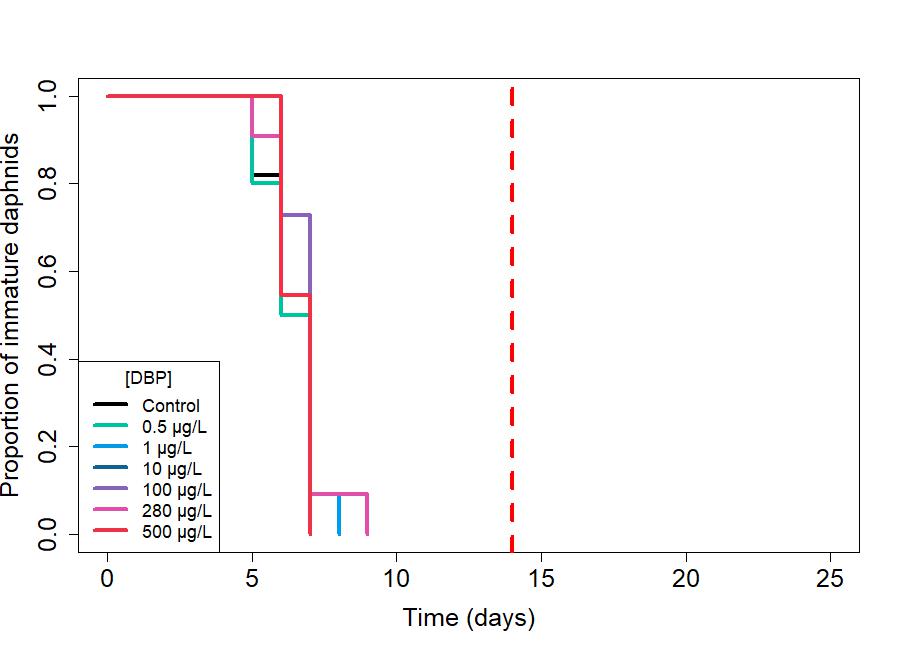

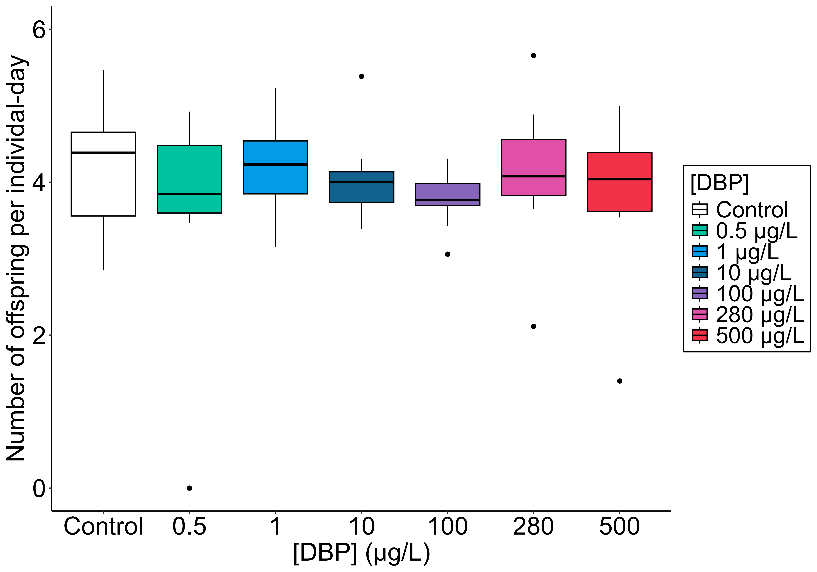

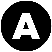

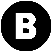

**Figure S4**. Reproduction of *D. magna* exposed to DBP under adult regime. (A) Time to sexual maturity of *D. magna* exposed to DBP under adult regime, expressed as the proportion of daphnids that have not yet laid their first brood among surviving individuals (n=11 for each concentration). *p ≤ 0.05 according to log-rank tests comparing each treatment to the control condition (p-value adjusted by a Bonferroni correction). (B) Fecundity of *D. magna* exposed to DBP under adult regime, expressed as number of offspring per individual-day as a function of concentration (n=11 for each concentration). *p ≤ 0.05 according to Wilcoxon tests comparing each treatment to the control condition (p adjusted by a Bonferroni correction). The horizontal line represents the median, the boxes range from the 25th to the 75th percentile and the whiskers extend to the last values within 1.5 times the inter-quartile range with black dots representing outliers.

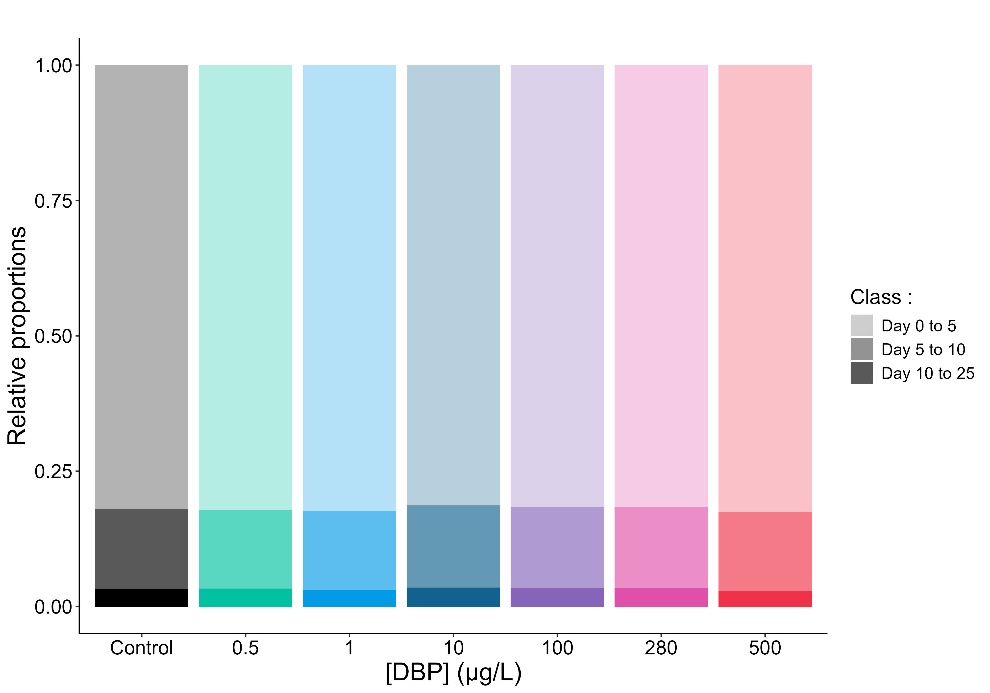

**Figure S5.** Stable age distribution of *D. magna* laboratory population for each DBP concentration under adult exposure. This stable age distribution was represented in three age classes (day 0 to 5, day 5 to 10 and day 10 to 25 days; faded colors).

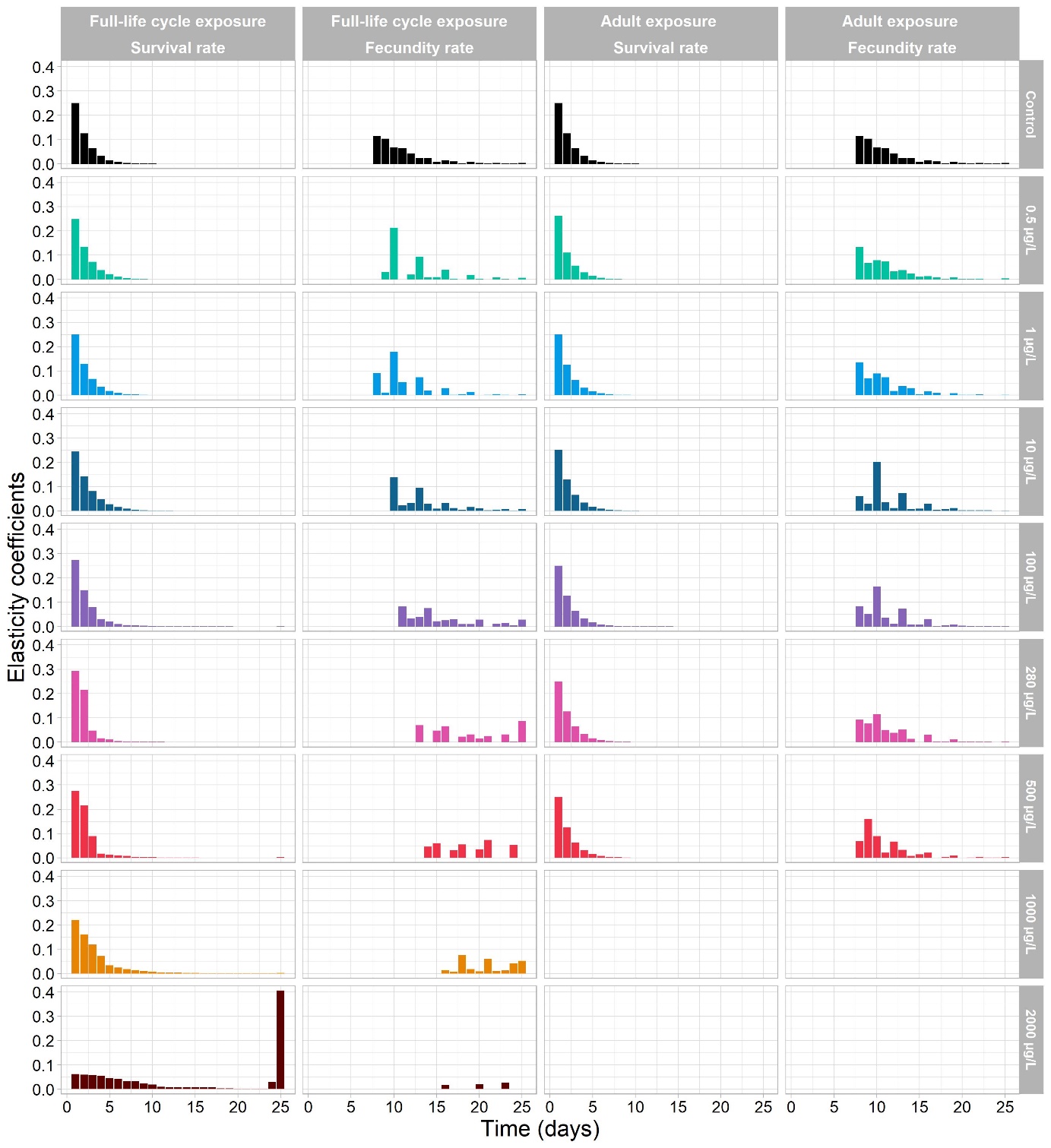

**Figure S6.** Elasticity analysis of *D. magna* populations exposed to DBP, reflecting contribution of survival and reproduction parameters to PRG determination. Survival and reproductive elasticity values for each regime and concentration.

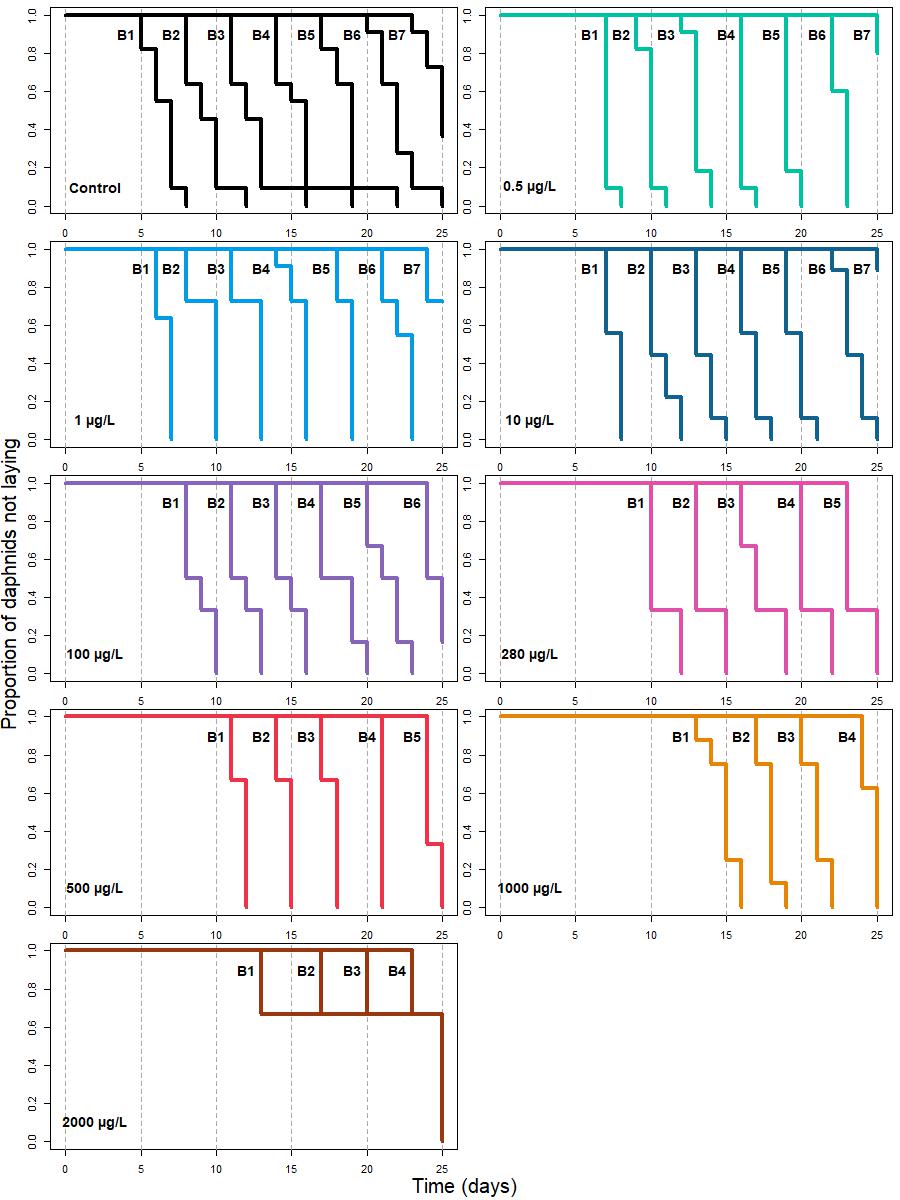

**Figure S7.** Time to successive broods in D. magna exposed to DBP under the full life cycle regime, expressed as the proportion of surviving individuals that have not yet laid their brood (n=11 for each concentration).

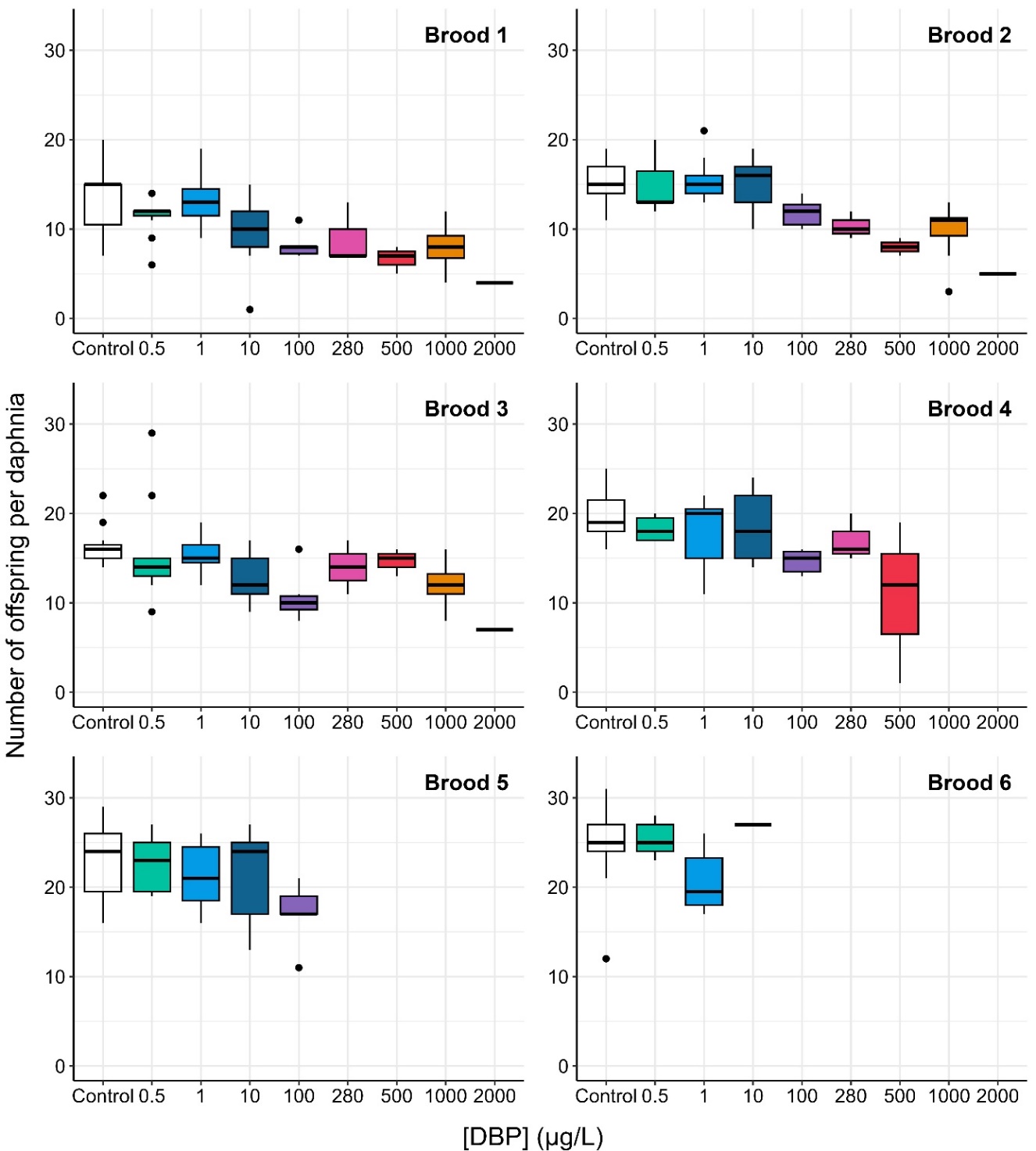

**Figure S8.** Fecundity of *D. magna* exposed to DBP under full life cycle regime, expressed as number of offspring per daphnid for each brood as a function of concentration (n=11 for each concentration). The horizontal line represents the median, the boxes range from the 25th to the 75th percentile and the whiskers extend to the last values within 1.5 times the inter-quartile range with black dots representing outliers.
